# Excessive visual crowding effects in developmental dyscalculia

**DOI:** 10.1101/2020.03.16.993972

**Authors:** Elisa Castaldi, Marco Turi, Sahawanatou Gassama, Manuela Piazza, Evelyn Eger

## Abstract

Visual crowding refers to the inability to identify objects when surrounded by other similar items. Crowding-like mechanisms are thought to play a key role in numerical perception by determining the sensory mechanisms through which ensembles are perceived. Enhanced visual crowding might hence prevent the normal development of a system involved in segregating and perceiving discrete numbers of items and ultimately the acquisition of more abstract numerical skills. Here, we investigated whether excessive crowding occurs in developmental dyscalculia (DD), a neurodevelopmental disorder characterized by difficulty in learning the most basic numerical and arithmetical concepts, and whether it is found independently of associated major reading and attentional difficulties. We measured spatial crowding in two groups of adult individuals with DD and control subjects. In separate experiments, participants were asked to discriminate the orientation of a Gabor patch either in isolation or under spatial crowding. Orientation discrimination thresholds were comparable across groups when stimuli were shown in isolation, yet they were much higher for the DD group with respect to the control group when the target was crowded by closely neighbouring flanking gratings. The difficulty in discriminating orientation (as reflected by the combination of accuracy and reaction times) in the DD compared to the control group persisted over several larger target flanker distances. Finally, we found that the degree of such spatial crowding correlated with impairments in mathematical abilities even when controlling for visual attention and reading skills. These results suggest that excessive crowding effects might be a characteristic of DD, independent of other associated neurodevelopmental disorders.

**Bullet points:**

- People with DD have difficulty learning about numbers and arithmetics.
- Perception of non-symbolic number seems to be modulated by visual crowding.
- Can stronger than normal crowding effects contribute to the origin of DD?
- We measured crowding with orientation discrimination tasks using Gabor gratings.
- Abnormal crowding characterizes DD independently of other developmental deficits.

## 1. Introduction

Developmental dyscalculia (DD) is a neurodevelopmental learning disability characterized by difficulty in learning about numbers and arithmetic, which manifests in children despite adequate neurological development, intellectual abilities and schooling opportunity (American Psychiatric Association, 2013). DD affects a wide range of mathematical abilities: DD individuals can have difficulties in understanding the meaning of numerical magnitudes and Arabic digits, in retrieving arithmetic facts from memory and in automatizing simple calculation procedures. Due to the variety of processes found to be often impaired, DD has been conceptualized as a multidimensional disorder: the difficulties in mathematical competence may emerge from the combination of weak domain-general functions (such as attention, memory and cognitive control) and domain-specific impairments in mastering numerical and arithmetical concepts (Fias et al., 2013; Fias, 2016; Iuculano, 2016).

Visuospatial working memory deficits occur in pure DD and these are stronger compared to profiles with associated dyslexia which are instead characterized by stronger verbal working memory deficits (Szűcs, 2016). Impairments in visuo-spatial attention and alertness have been reported in DD individuals, despite not meeting the criteria for attention-deficit/hyperactivity disorder (ADHD) (Askenazi and Henik, 2010). During numerical processing, calculation and math problem solving, these domain-general functions are thought to interact with a specific ‘core knowledge’ of magnitude, foundational for mathematical competence. Since very early in life, humans can perceive changes in the number of objects in an image (Brannon et al., 2004; Izard et al., 2009; Libertus et al., 2014; de Hevia, 2016; 2017), an ability that is thought to be based on our ‘number sense’ (Dehaene, 1997). During development, numerical magnitudes, initially experienced in their non-symbolic format, are thought to be mapped onto their symbolic counterpart, setting the base for formal arithmetical learning (Piazza, 2010). In line with this view, the ability to discriminate between non-symbolic numerical quantities (also known as ‘numerical acuity’) was found to be predictive of arithmetical skills in the neurotypical population (Halberda et al., 2008; Piazza et al., 2013; Chen and Li, 2014; Anobile et al., 2016a, 2018) and to refine with education (Piazza et al., 2013). Importantly, these basic numerical abilities were found to be disproportionally impaired in DD individuals: they tend to be slower and/or less accurate than their age-matched peers when briefly shown with two sets of dots or with two digits and asked to choose the numerically larger one or when tested with estimation tasks requiring mapping numbers across formats (Butterworth, 2005; Rousselle and Noël, 2007; Iuculano et al., 2008; Piazza et al., 2010; Butterworth, 2010; Mejias et al., 2012). In line with these observations, some of the most influential models of DD attributed the origin of the numerical and mathematical difficulties to a specific deficit in the core representation of quantity (Dehaene et al., 2003; Butterworth, 2005, 2010).

If DD originates from, among other factors, a weak number sense, then it is important to investigate whether the visual mechanisms supporting the extraction of numerosity from an image are impaired. To date, the exact visual mechanisms mediating numerosity perception are still a matter of debate. Some authors suggested that number is sensed directly through dedicated “number detectors” (Burr and Ross, 2008; Ross, 2010), while others proposed instead that number is perceived through mechanisms related to texture-density processing (Dakin et al., 2011; Morgan et al., 2014). Importantly, behavioural studies in individuals without math difficulties have provided evidence that the mechanism through which ensembles are perceived can change depending on how cluttered items are (Anobile et al., 2016b): Anobile et al. (2013) measured discrimination thresholds as a function of dot numerosity and found that, for relatively sparse arrays of smaller numerosites, the thresholds followed Weber’s law (remained constant across numbers), however, for larger numerosities when sets became highly cluttered, the discrimination thresholds decreased steadily with numerosity, following a square-root law. The fact that discrimination thresholds obeyed different psychophysical laws suggested that visual arrays can be processed by two independent perceptual systems: one related to number perception, recruited when viewing sufficiently sparse arrays, and the other related to texture-density perception, recruited when individual items are too cluttered to be clearly segregated (Anobile et al., 2015, 2016b; Burr et al., 2017). Interestingly, only the ability to discriminate the numerosity of sparse, but not of cluttered arrays, was found to predict numerical skills in typically developing children (Anobile et al., 2016a), suggesting that only the functionality of the system mediating number perception, and not of the one mediating texture-density perception, may be relevant for the development of formal mathematical abilities. When studying the variation of discrimination thresholds as a function of dot numerosity, Anobile et al. (2015) observed that the switching point between the two psychophysical regimes depended on the number of dots per visual degree and on the eccentricity at which the arrays were shown, while being unrelated to the individual items’ size. These observations suggested that the recruitment of one of the two systems supporting ensemble perception might be regulated by mechanisms related to visual crowding, a process which sets limits to our ability to identify, locate and count objects when they are cluttered together (for reviews see: Levi, 2008; Pelli, 2008; Pelli and Tillman, 2008; Whitney and Levi, 2011). Crowding degrades visual perception, making individual items appear jumbled together with the surrounding objects (Pelli et al., 2004). When multiple nearby items are merged into a single percept, numerosity might undergo underestimation: Valsecchi et al. (2013) showed that the perceived numerosity of arrays presented in the periphery was reduced compared to central viewing, a phenomenon that could be simulated by a texture synthesis algorithm inducing texture formation of cluttered items in random dot arrays (Balas, 2016). The perceived numerosity was found to decrease for smaller inter-item distance, and this effect could not be accounted by blurring in peripheral vision nor by misperception of stimulus size, leading Valsecchi et al. (2013) to attribute the underestimation of peripherally viewed arrays to visual crowding (however see: Chakravarthi and Bertamini, 2020, for an alternative interpretation of underestimation based on clustering rather than crowding). Interestingly, crowding may also lead to perception of additional elements in the scene: using a change detection task, Greenwood et al. (2010) showed that crowing induced participants to perceive a target noise or even an inexistent target (blank space) to be oriented as the flanking gratings.

If mastering non-symbolic numerical quantities sets the basis for the development of formal mathematical competence and if numerosity perception is limited by crowding, could stronger than normal crowding mechanisms contribute to the origin of DD? By hampering efficient individual items’ identification, abnormal crowding mechanisms might impair ensemble perception leading to the recruitment of texture-density mechanisms even for relatively sparse ensembles. This might then induce a cascade of events: because items might appear too cluttered to be clearly segregated, the representation of numerosity might be imprecise and understanding the meaning of symbolic numbers more difficult, posing challenges to the development of efficient symbolic mathematical skills.

The effect of crowding on the development of numerical cognition has not been investigated so far. Crowding has been classically studied using letter stimuli (Flom et al., 1963; Bouma, 1970; Toet and Levi, 1992) for its effect on reading: we can only read letters within our central “uncrowded” window while letters in the periphery will be undistinguishable. Our reading rate is limited by crowding, it depends on the observer’s critical spacing (the smallest distance between items that avoids crowding) and on the spacing between the viewed letters (Legge et al., 2001; Pelli et al., 2007; Yu et al., 2007). There occurs a developmental expansion of the size of this uncrowded window from the 3rd grade to adulthood which parallels reading speed in normal readers (Bondarko and Semenov, 2005; Kwon et al., 2008). Interestingly, there is evidence that some individuals with developmental dyslexia, a specific learning disability affecting reading skills, have increased crowding with respect to proficient readers (Bouma, 1970; Spinelli et al., 2002; Martelli et al., 2009; Moores et al., 2011; Perea et al., 2012; Zorzi et al., 2012; Callens et al., 2013; Moll and Jones, 2013; Cassim et al., 2014; Montani et al., 2015; for a review see: Gori and Facoetti, 2015). More recently it has been proposed that crowding might contribute to the origin of developmental dyslexia (Spinelli et al., 2002; Gori and Facoetti, 2015), although it cannot fully account for it: dyslexics’ reading rate still remains slower than proficient readers even after compensating for crowding (Martelli et al., 2009).

Given the reviewed evidence for a crucial contribution of crowding-like mechanisms to the processing of visual numerosity, in the current study we investigated whether excessive crowding is present in DD. Importantly, given that DD is often found in comorbidity with developmental dyslexia (Rubinsten, 2009; Wilson et al., 2015), we tested whether abnormal crowding mechanisms are present in DD independently of associated reading disorders. Moreover, given that DD is also often associated with ADHD (Rubinsten, 2009) and considering that one of the most influential model of crowding attributed the occurrence of this phenomenon to limited attentional resources (He et al., 1996), in the current study we also evaluated crowding in DD independently of major visual attentional deficits, potentially indicative of ADHD. To this aim, we measured visual crowding effects in DD individuals and compared their performance against a group of matched controls. In order to investigate visual crowding independently of non-symbolic number comparison abilities, we used stimuli and tasks that were not related to number processing and asked participants to judge the orientation of a Gabor patch shown in isolation or surrounded by flankers. We further measured participants’ ability to discriminate sparse and dense non-symbolic ensembles, hypothesising that DD subjects would specifically fail in the former, suggesting a specific fragility of the number system. Finally, we explored the relation between visual crowding, numerical and mathematical abilities, and we speculate on the possible contribution of crowding mechanisms in the development of numerical cognition.

## 2. Material and methods

### 2.1 Subjects

Seventeen adults without mathematical impairment and seventeen adults with mathematical impairment were included in the study. Participants without mathematical impairment were recruited through a diffusion list provided by the CNRS, primarily directed to cognitive psychology students, but open also to students from other faculties.

Contacts with participants with mathematical impairment were provided either by our speech therapist and neuropsychologist collaborators or obtained through advertisements on social media and in universities. The advertisement encouraged people with mathematical difficulties to fill in an online screening questionnaire. In addition to collecting general information (such as age and schooling level), the first part of the questionnaire investigated whether the individual had received a diagnosis of dyscalculia or neurological disorders. The second part of the questionnaire explored whether the claimed math difficulties were sufficiently strong to impact the individual’s everyday life (for example when dealing with money or quantities in social environments and everyday activities), or to impact the ability to perform some basic numerical tasks (such as counting or reading/writing numerals or solving simple mathematical operations without using fingers or a calculator).

To be included in the experiment, all participants were required to (a) be between 18 and 50 years old, (b) present no neurological disorder, and (c) have completed at least secondary level education. Furthermore, participants that were included in the math impaired group needed to either have been clinically diagnosed with dyscalculia by a neuropsychologist or speech therapist or have claimed major difficulties when dealing with numbers according to the questionnaire. Participants fulfilling these criteria were contacted to participate in an extensive neuropsychological assessment, where measures of verbal and non-verbal intelligence, verbal and visuospatial working memory, visual attention, reading abilities, inhibitory skills and mathematical performance were obtained. In a separate session held in a different day, participants performed a series of psychophysical experiments.

Two subjects included in the math impaired group dropped from the study: one subject who initially showed interest in participating in the study was not available for the proposed testing sessions and not further contactable afterward. The other subject underwent the neuropsychological assessment but was never available for the second testing session requiring participants to perform the psychophysical tests.

To define the final DD and control groups, we performed an additional selection based on the results of the neuropsychological assessment. Specifically, we z-scored the participants’ results to the math tests: first, we calculated the mean and standard deviation of the scores obtained by the participants without math difficulty in each test, then we used these values for normalization, i.e. we subtracted the mean of the group without math difficulty from the score of each participant (including DD) and then divided it by the standard deviation of the participants without math difficulty. Mathematical performance was considered below the normal level if z-scores calculated from either accuracy or reaction time in two (of a total of four) or more math tests exceeded the average z-scores of the non-math-impaired group by more than 2 standard deviations. All DD participants exceeded this cut off. Two participants in the non-math-impaired group exceeded this cut off and where therefore discarded. The same procedure was applied to the accuracy and reading speed of a reading test and of a visual search test (see below for test description) in order to identify DD subjects who also had major associated reading or attentional deficits, potentially reflecting associated dyslexia or ADHD disorders. Two DD subjects exceeded the cut-off for reading abilities (one for number of errors, the other for speed) and other two exceeded the cut-off for the number of errors in the visual search test.

Overall, fifteen participants in the control group (age 31±10, 8 females) and fifteen participants in the DD group (age 27±11, 10 females), four of which with associated reading or attention difficulties, were included in the study. All participants had normal or corrected to normal visual acuity.

Written informed consent was obtained from all participants in accordance with the Declaration of Helsinki, and the study was approved by the research ethics committee of University Paris-Saclay.

### 2.2 Neuropsychological Assessment

Prior to the psychophysical experiments, subjects underwent neuropsychological assessment. We selected the subtests Similarities and Matrix Reasoning from the Wechsler Adult Intelligence Scale IV edition (WAIS-IV) as a measure of verbal and non-verbal IQ. As a measure of verbal working memory, we selected the digit span subtest from WAIS-IV, while the Corsi Block Tapping test was used to measure visuospatial working memory.

Reading abilities were assessed with the “Alouette” (Lefavrais, 1967), the most widely-used French reading test. It involves reading aloud a brief text composed of grammatically plausible sentences, including existing regular and irregular words, without a clear overall meaning. The time needed to read the text and the number of errors made were measured.

To measure inhibitory skills, the Stroop-Victoria test adapted for francophone subjects (Bayard et al., 2009) was administered. Participants were required to pronounce as quickly as possible the color of the ink of a series of filled circles, of a list of words (‘mais’, ‘pour’, ‘donc’, ‘quand’, meaning ‘but’, ‘for’, ‘so’, ‘when’) and of a list of color words (‘jaune’, ‘rouge’, ‘vert’, ‘bleu’, meaning ‘yellow’, ‘red’, ‘green’, ‘blue’). For the color words, the color of the ink was always incongruent with the meaning (for example ‘rouge’, meaning red, written in blue). The interference index was calculated by dividing the time needed to perform the task with the color words by the time needed to name the color of circles.

As a measure of visual attention, visual search performance was measured with the Bells test. Participants were shown a paper sheet containing black silhouettes of different objects. They were required to identify and cross with a pen all the bells embedded in the sheet. Time recording was stopped when the participant considered all the bells crossed and the number of omitted bells was recorded.

To evaluate mathematical abilities, participants were tested with several subtests of the French battery TEDI Math Grands (Noël and Grégoire, 2015). This computerized battery measures the individual’s performance over a range of tests targeting various basic numerical abilities. Subjects were required to: 1) estimate the number of briefly presented dots within the small numerosity range (1-6 items); 2) compare two single-digit Arabic numerals; 3) mentally perform single-digit multiplications and subtractions. The software collected the participants’ accuracies and reaction times for most of these tests, with the exception of the test measuring the ability to estimate numerosities for which only the accuracy was recorded.

We calculated standard scores for the IQ subtests (Similarities and Matrix reasoning), for the verbal and visuospatial working memory and for inhibition referring to standardized norms for adults. For the TEDI-MATH we analyzed the number of correct responses and, when recorded, the reaction time (in ms) needed to respond. Given that reaction time and accuracy can often inversely trade off with each other, we reduced the number of variables by calculating the inverse efficiency score (IES, Collins et al., 2017), corresponding to the reaction time (RT) divided by the proportion of correct responses. From the TEDI-MATH results, we computed: 1) IES Digits obtained from the results of the single-digit Arabic numeral comparison test; 2) IES Calculation – obtained by averaging together the results from the multiplication and subtraction tests and computing the IES from the combined measure; 3) IES Math – obtained by averaging the IES Digits and IES Calculation as index of general math ability.

Independent sample t-tests were performed to evaluate differences across groups. These tests were applied to either the standardized test scores described (for the IQ, memory and inhibition tests) or to the raw scores in the cases where the norms did not cover the adult age range (in the case of the math, reading and visual search tests).

### 2.3 Psychophysical experiments

All visual stimuli used in the psychophysical experiment were viewed binocularly from approximately 60 cm in a dimly lit room, displayed on a 15-inch Laptop (HP) LCD monitor with 1600×900 resolution at refresh rate of 60 Hz. Stimuli were generated and presented under Matlab using PsychToolbox routines (Brainard, 1997).

#### 2.3.1 Visual crowding: Experiments 1-3

Stimuli were oriented Gaussian-windowed sinusoidal gratings (carrier frequency 1 cycle/visual degree, the gaussian window around the Gabors had a standard deviation of 0.85°, Michelson contrast of 0.9%) presented on a gray background. Stimuli were displayed randomly to the left or right of the central fixation point at 10° of eccentricity (Figure 1). Subjects were asked to maintain central fixation and to judge whether the external half of the grating (with respect to the screen center) was tilted up or down of horizontal by pressing either the up or down arrows on the keyboard. At the beginning of each trial, a central fixation point was shown for 1500 ms, then the stimuli were briefly presented for 33 ms. Participants were instructed to provide their response in the shortest time possible. In Experiment 1, the oriented grating (Figure 1A) was presented in isolation, whereas in Experiment 2 (Figure 1B), the target grating was presented simultaneously with two other gratings (identical features, but with fixed horizontal orientation) that served as distractors (flankers). The two flankers vertically surrounded the target, at a fixed and very close center-center distance (0.85°). In Experiment 2, participants were asked to report the orientation of the central grating (target), ignoring the two flankers.

**Figure 1.**
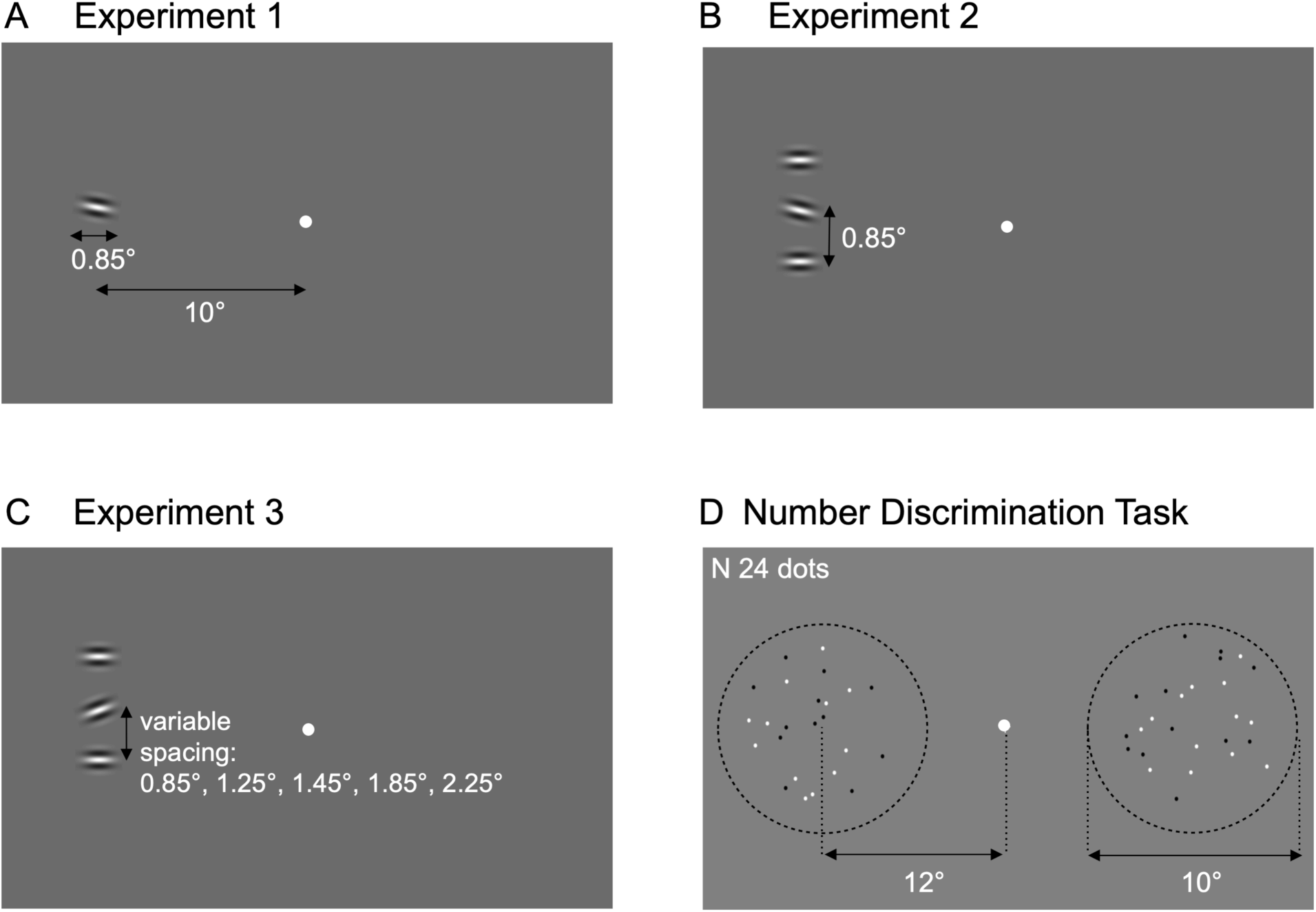
Examples of stimuli used in the visual crowding and number discrimination experiments. (A) In Experiment 1, a single oriented Gabor grating was briefly shown (33ms) at 10° of eccentricity either to the left or right of the central fixation point. Participants were asked to judge the grating’s orientation with respect to the imaginary horizontal axis and to press the up or down arrow key (the correct response in this trial is ‘up’). (B) In Experiment 2, three oriented gratings were shown at a very close distance and the participants’ task was to judge the orientation of the central one, ignoring the two flankers. (C) Stimuli and task in Experiment 3 were the same as in Experiment 2, but the target-flanker distance was varied, to modulate the strength of the crowding effect (the larger the distance, the weaker the crowding effect). (D) In the number discrimination task, two arrays of dots were briefly (250 ms) and simultaneously presented. Subjects were asked to indicate which side of the screen contained more dots.

In both Experiment 1 and 2, the target orientation was adaptively changed according to the subject’s responses following a QUEST algorithm (Watson and Pelli, 1983). Participants performed one session of 60 trials for each experiment. Responses faster than 150 ms or longer than two standard deviations with respect to each participant’s average reaction time in each condition were discarded from the analysis. For each participant, the proportion of correct responses against the target orientation angle was fitted by a cumulative Gaussian function. The orientation angle needed for the subject to score 75% correct defined the participant’s orientation threshold. To verify the reliability of the estimated thresholds, we calculated the goodness of fit for each subject and experiment and set a minimum limit at R^2^ =0.75. The quality of the fit did not meet this criterion for the data obtained from one DD participants in Experiment 1 and two DD participants in Experiment 2, confirming their verbally reported difficulty in performing these tasks. Given that a reliable estimate of the orientation discrimination threshold in the condition tested in Experiment 2 was needed to perform Experiment 3 (see below), data from these three subjects was discarded from the analyses of Experiment 1-3. Orientation discrimination thresholds and reaction times measured in Experiment 1 and 2 were entered into a repeated measures ANOVA with crowding condition (2 levels: uncrowded/crowded) and group (2 levels: control/DD) as within- and between-subject factors, respectively.

Similar to Experiment 2, in Experiment 3 participants were presented with three vertically positioned oriented gratings and asked to evaluate the orientation of the central target (Figure 1C). The central target orientation was fixed at the participant’s orientation discrimination threshold, estimated from Experiment 2. The target-flanker distance was varied at every trial to measure the spatial extent of crowding (i.e. the critical spacing). Five target-flanker distances were tested: 0.85° (same as Experiment 2), 1.25°, 1.45°, 1.85° and 2.25°. Trials in Experiment 3 were presented using the method of constant stimuli: participants performed 20 trials for each of the 5 target-flanker distances and 2 presentation sides (left and right of the fixation point), for a total of 200 trials. As for Experiment 1 and 2, responses faster than 150 ms or longer than two standard deviations with respect to each participant’s average reaction time were discarded from the analysis. Response accuracies were entered into a repeated measures ANOVA with target-flanker distance (5 levels: the 5 target-flanker distances) and group (2 levels: control/DD) as the within- and between-subject factors respectively. Furthermore, we computed the critical spacing, defined as the target-flanker distance at which the response accuracy reached the plateau value of 90%, following the procedure used by (Freyberg et al., 2016). To this end, we plotted the accuracy as a function of the target-flanker distance and fitted the data with an exponential curve using the following equation:

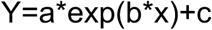

where x refers to the target-flanker distance, with the constraints that the parameters a and b had to be negative, and c had to fall between 0.5 and 1. As starting values for the parameters a, b, and c we chose -0.2, -2, and 0.95 respectively. This ensured that the percent accuracy increased with target-flanker distance and plateaued to a value between 50% and 100% accuracy. The critical spacing was therefore calculated as:

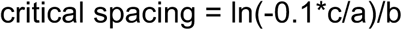

and compared across groups by means of independent sample t-test.

Reaction times were also recorded and combined with response accuracies to calculate the inverse efficacy score (IES). Reaction times and IES were entered into ANOVA with target-flanker distance (5 levels: the 5 target-flanker distances) and group (2 levels: control/DD) as the within- and between-subject factors, respectively.

Similarly to (Cassim et al., 2014), we calculated a crowding index defined as the difference between IES at the smallest (0.85°) and the largest (2.25°) target-flanker distance. This measure reflects the accuracy-normalized reaction time cost of performing the orientation discrimination task in the stronger crowding compared to weaker crowding condition.

The statistical analyses which resulted significant when testing the whole group of DD subjects who successfully performed Experiment 1-3 (twelve subjects), were replicated using a reduced group which did not include the four DD subjects who had associated reading or attentional difficulties.

#### 2.3.2 Non-symbolic Number Discrimination Task

Stimuli consisted of array of dots, half white and half black, presented on a mid-gray background, so that luminance was not a cue for number. Two arrays of dots were simultaneously and briefly (250 ms) presented on either side of the central fixation point, at 12° of eccentricity (Figure 1D). Participants performed a comparison task indicating the side of the screen with the more numerous array. In separate sessions, the numerosity of the probe array (randomly shown to the left or to the right of the fixation point) was fixed at 24 (N24, sparse array) or at 64 (N64, dense array) dots, while the numerosity of the test array adaptively changed, according to the participant’s responses, following the QUEST algorithm. Dots were 0.1° diameter large and constrained to fall within a virtual circle of 10° diameter (yielding a density of 0.3 and 0.82 dots/deg^2^ for N24 and N64, respectively). The minimal inter-dot distance was 0.3°.

Participants performed two separate sessions of 35 trials each, testing the two numerosities, with half the participants starting with N24 and the other half with N64. The proportion of trials where the test appeared more numerous than the probe was plotted against the logarithm of the test numerosities and fitted with a cumulative Gaussian function. The mean of this function provided an estimate of the point of subjective equality (PSE, a measure of perceived numerosity), while the standard deviation was used to estimate the precision (i.e., the just-noticeable difference, JND), which was divided by the point of subjective equality to estimate the Weber fraction. Weber fractions were entered into an ANOVA with numerosity (2 levels: N24 and N64) and group (2 levels: control/DD) as within and between subjects’ factors respectively. All fifteen DD subjects successfully performed the experiment and were entered in the analysis.

## 3. Results

### 3.1 Neuropsychological assessment

In the interview conducted during the neuropsychological assessment, all participants confirmed being free of neurological disorders and having had access to appropriate education during school-age. Among the participants included in the DD group, four had received formal diagnosis of dyscalculia during childhood and the others confirmed having always had difficulties whenever dealing with numbers and quantities and major problems in acquiring mathematical skills since the early school years. All subjects claimed that the mathematical difficulties persisted over years. Furthermore, 7 participants out of 15 reported having at least one relative with difficulty in mathematics, reading, writing, or orthography.

The DD and control group were not significantly different in age, verbal and non-verbal IQ, reading accuracy, inhibitory control, as measured by the Color-Stroop test, and visual search performance, as measured by the Bells test, (all p-values>0.05, see Table 1 for descriptive statistics and tests across groups). The DD and control groups significantly differed in reading speed (t(27)=2.47, p=0.02), verbal (t(28)=-2.59, p=0.01) and visuo-spatial working memory (t(28)=-3.27, p=0.002), and most of the numerical and arithmetical tests. Specifically, the DD group was slower when comparing digits (t(28)=3.97, p=0.0004), performing mental multiplication (t(28)=4.34, p=0.0002) and subtraction (t(28)=4.79, p=0.00005). Accuracy for mental multiplication and subtraction was also significantly lower with respect to the control group (for multiplication: t(28)=-5.05, p=0.00002; for subtraction t(28)=-2.18, p=0.04). IES for digit comparison (t(28)=4.03, p=0.0004), calculation (t(28)=5.20, p=0.00002) and general math (t(28)=2.77, p=0.009) significantly differed across groups.

**Table 1.**

|  | Control group<br>(N=15) | Dyscalculic group<br>(N=15) | Statistical<br>analysis |
| --- | --- | --- | --- |
|  | Mean (STD) | Mean (STD) | t-value |
| Age | 31 (10) | 27 (11) | -0.93 |
| IQ |  |  |  |
| Similarities | 13 (3) | 13 (2) | 0.42 |
| Matrices | 12 (3) | 10 (3) | -1.83 |
| Reading Ability |  |  |  |
| Time (seconds) | 89 (14) | 106 (22) | 2.47 * |
| N errors | 3 (3) | 4 (3) | 1.43 |
| Working memory |  |  |  |
| Verbal (Digit span) | 12 (3) | 9 (3) | -2.59 * |
| Visuospatial (Corsi) | 13 (2) | 10 (2) | -3.27 ** |
| Inhibition |  |  |  |
| Color Stroop Score | 12 (2) | 11 (4) | -1.15 |
| Visuo-Spatial attention |  |  |  |
| Time (seconds) | 105 (45) | 112 (37) | 0.51 |
| N omissions | 1 (1) | 2 (3) | 0.53 |
| Numerical skills /<br>Arithmetics TEDI-MATH<br>(no of items) |  |  |  |
| Small numerosity<br>estimation (36) | 33 (4) | 32 (3) | -0.75 |
| Digit Comparison<br>Accuracy (48) | 46 (1) | 47 (2) | 1.04 |
| Reaction Time (ms) | 558 (51) | 739 (168) | 3.97 ** |
| IES Digit (ms) | 578 (54) | 754 (159) | 4.03 ** |
| Multiplication<br>Accuracy (20) | 18 (2) | 15 (2) | -5.05 ** |
| Reaction Time (ms) | 1681 (403) | 3617 (1681) | 4.34 ** |
| Subtraction<br>Accuracy (20) | 19 (1) | 18 (2) | -2.18 * |
| Reaction Time (ms) | 1572 (333) | 3307 (1362) | 4.79 ** |
| Calculation (x and -)<br>IES Calculation (ms) | 3481 (727) | 8365 (3559) | 5.20 ** |
| IES General Math (ms) | 1166 (201) | 2007 (1156) | 2.77 ** |
| DD differs significantly from controls at: * $p < 0.05$, ** $p < 0.01$ | | | |

### 3.2 Visual Crowding

One participant from the DD group did not succeed in completing Experiment 1, which required to judge the orientation of one isolated grating. The participant claimed that the difficulty was not due to discriminating the orientation of each individual grating, but to associating a pair of orientations (each one the mirror image of the other with respect to a vertical imaginary symmetry axis) with the corresponding response (either up or down arrows). To check that the participant did not have a general orientation discrimination impairment we measured the ability to discriminate the orientation of a central grating around +45 or –45 degrees (for details on the stimuli and procedure see: Castaldi et al 2018a). The participant showed very good thresholds for both orientations (3 and 1 visual degrees for +45 and –45 degrees respectively),comparable to typical subjects’ average discrimination threshold of 4.4±0.7 (typical values taken from: Castaldi et al., 2018a). Two other subjects from the DD group, while easily performing Experiment 1, failed to complete Experiment 2, in which the target grating was surrounded by two flankers placed in close proximity. The participants claimed that the task was too difficult because they could not distinguish the central target from the flankers. These qualitative verbal reports were confirmed by the poor data quality which did not allow the fitting to provide reliable thresholds (goodness of fit R^2^ lower than 0.15 for both subjects, see methods). As a consequence, it was not possible to test these three subjects with Experiment 3 for which a measure of the participant’s orientation threshold at the closest target-flanker distance (estimated in Experiment 2) was needed. These subjects were therefore excluded also from Experiment 1 and the results described below were obtained from the remaining twelve subjects included in the DD group.

Results from Experiment 1 and 2 are shown in Figure 2. Figure 2A shows the psychometric curves of two representative single subjects from the control (Figure 2A) and DD (Figure 2B) group in Experiment 1 (uncrowded condition, hatched curves) and Experiment 2 (crowded condition, solid curves). The rightward shift of the solid psychometric curves on the x-axis indicates that the orientation discrimination thresholds increased under crowding for both groups. The orientation discrimination thresholds measured in the two groups and crowding conditions are shown in Figure 2C. Orientation discrimination thresholds were similar across groups in the uncrowded condition (2.57°±1.19° for the control group and 3.53°±2.25° for the DD group) and increased in the crowded condition in both groups. However, the presence of flankers surrounding the target induced a much larger increase in the orientation discrimination thresholds in the DD group with respect to the control group (8.12° ± 2.37° for the control group and 12.48° ± 5.96° for the DD group). To statistically test for these differences, we entered the orientation discrimination thresholds into a repeated measures ANOVA with crowding condition (2 levels: uncrowded/crowded) and group (2 levels: DD/controls) as within- and between-subject factors, respectively. The significant interaction between crowding condition and group (F(1,25)=5.01, p=0.03, ηp^2^=0.16) and the post-hoc tests confirmed that although the orientation discrimination thresholds under crowding significantly increased both in the DD (t(11)=-5.89, p<10^−5^, Cohen’s *d*=1.69) and in the control group (t_(14)_=-8.97, p<10^−5^, Cohen’s *d*=2.32) with respect to the uncrowded condition, this increase was significantly stronger for the DD group with respect to the control group (t_(25)_=-2.59, p=0.016, Cohen’s *d*=0.96). Importantly, this difference could not be attributed to an overall poorer orientation discrimination ability in DD, as the orientation discrimination thresholds in the uncrowded condition were not statistically different across groups (t_(25)_=-1.41, p=0.17, Cohen’s *d*=0.55).

**Figure 2.**
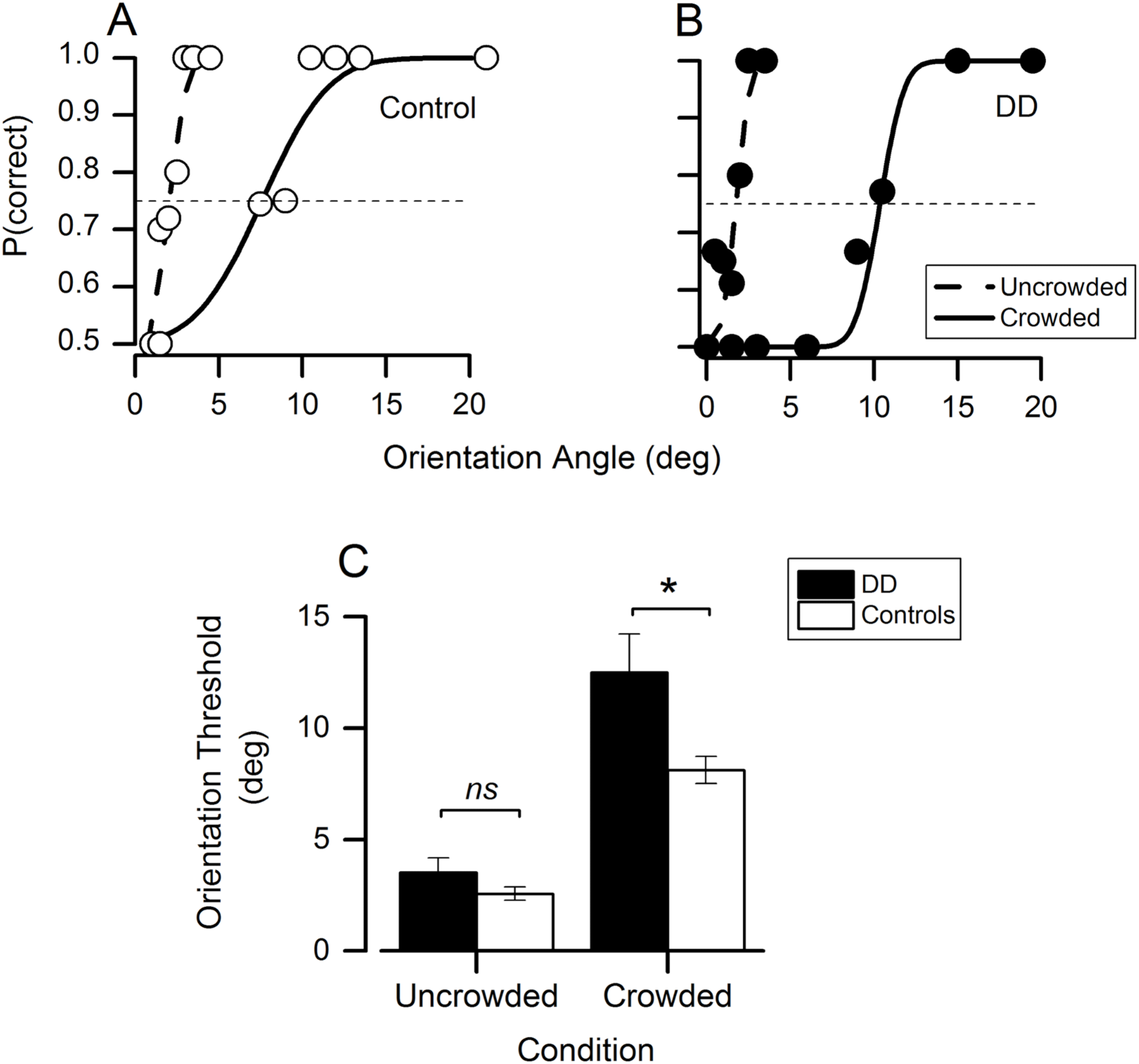
Effect of visual crowding on orientation discrimination thresholds. Psychometric functions for two subjects from the control (A) and the DD (B) group. The proportion of correct responses is plotted as a function of the orientation angle with respect to the horizontal (in visual degrees) for the uncrowded (hatched curve, Experiment 1) and crowded (solid curve, Experiment 2) conditions. Orientation thresholds were measured at the point in which the proportion of correct responses reached 75% (dashed lines). (C) Average orientation discrimination thresholds in the DD (black bars) and control (white bars) groups for the two crowding conditions. The addition of flankers affected orientation discrimination performance in both groups, but this effect was stronger in the DD group with respect to the control group.

Reaction times of the DD group were comparable to the control group in the uncrowded condition (RTs control group=0.81±0.15, RTs DD group= 0.97±0.25) and visual crowding slowed down reaction times in both groups (RTs control group=0.93±0.21, RTs DD group=1.05± 0.17). Repeated measure ANOVA revealed a significant main effect of crowding condition (F(1, 25)=8.97, p=0.006, ηp2=0.26), but no significant main effect of group (F(1,25) = 3.88, p = 0.06, ηp2=0.13), nor significant interaction between group and crowding condition (F(1, 25)=0.15, p=0.70, ηp2=0.006), suggesting that visual crowding slowed down reaction times in the two groups to the same extent.

In sum, orientation discrimination thresholds in peripheral vision were comparable across groups when stimuli were presented in isolation, and visual crowding affected performance in both groups, as reflected by both the higher orientation discrimination threshold and the longer reaction times in the crowding condition. However, visual crowding caused a stronger increase in the orientation discrimination thresholds in the DD group with respect to the control group.

In Experiment 3 we measured the spatial extent of the visual crowding effect in the two groups, by fixing the target orientation at each individual subject’s threshold, measured in Experiment 2, and varying the target-flanker distance. The orientation discrimination performance was expected to progressively improve with larger target-flanker distances, as the flankers should progressively fall outside the crowding window and stop interfering with the central target perception. Figure 3A shows that indeed the proportion of correct responses increased with larger target-flanker distances in both groups. The proportion of correct responses was entered into a repeated-measures ANOVA, with target-flanker distance (5 levels) and group (2 levels) as the within- and between-subject factors, respectively. The proportion of correct responses started at 0.75 for the shortest target-flanker distance (as expected given that this distance corresponded to the one at which the orientation threshold was estimated in Experiment 2), and increased with larger target-flanker distances in both groups (significant main effect of target-flanker distance: F_(4,44)_=53.20, p<10^−5^, ηp^2^=0.89), up to 0.9. The magnitude of this effect was comparable across the two groups (no significant main effect of group: F_(1,25)_=0.14, p=0.71, ηp^2^=0.01; no interaction between group and target–flanker distance: F_(4,44)_=1.97, p= 0.12, ηp^2^=0.14). The critical spacing, although on average slightly higher in the DD with respect to the control group (DD group: 1.78°±0.88°, control group: 1.57°±0.79°), was not significantly different (t_(25)_=0.64 p=0.52, Cohen’s *d*=0.25).

**Figure 3.**
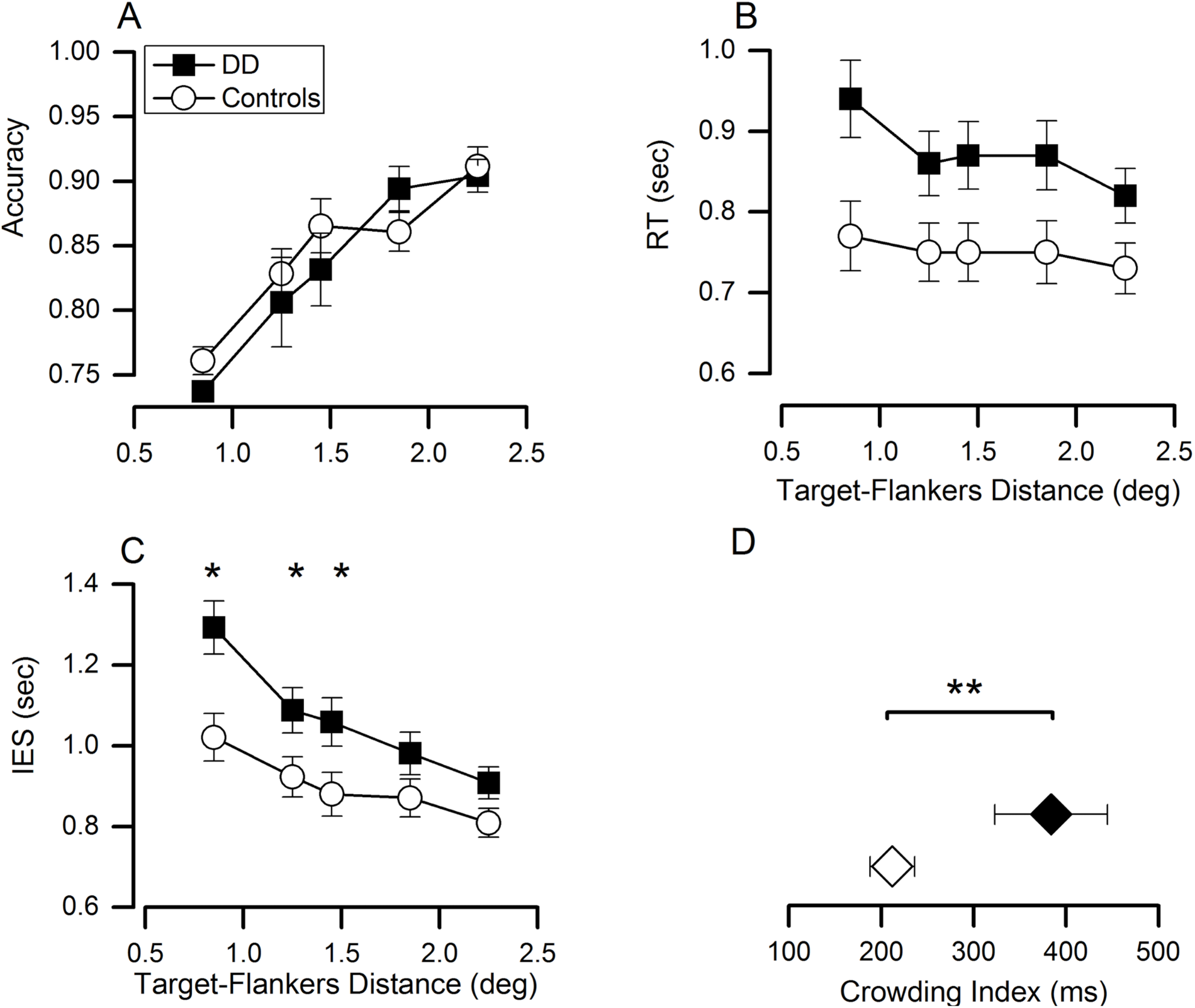
Spatial extent of the visual crowding effect. (A) The proportion of correct responses is plotted as a function of the target-flanker distance for the DD (black symbols) and the control (white symbols) group. As expected, response accuracy increased with larger target-flanker distance. Reaction times (B) and IES (C) decreased with larger target-flanker distance in both groups. IES was significantly higher in DD with respect to the control group, especially for the closest target-flanker distances. The crowding index (D) was higher in the DD with respect to the control group.

Reaction times (Figure 3B) and IES (Figure 3C) decreased as a function of the target-flanker distance, meaning that responses speeded up and the orientation discrimination performance improved as the flankers were displayed farther away from the target in both groups.

A repeated measures ANOVA confirmed that reaction times significantly decreased with target-flankers distance in both groups (significant main effect of target-flanker distance: F_(4,100)_=8.95, p<10^−5^, ηp^2^=0.26), however the DD group was overall slower with respect to the control group (significant main effect of group: F_(1,25)_=5.31, p=0.03, ηp^2^=0.18, no significant interaction between group and target–flanker distance: F_(4,100)_=2.21, p=0.07, ηp^2^=0.08).

These results suggest that in order to perform the orientation discrimination task with response accuracy comparable to the control group, the DD group needed more time. The IES scores decreased with target-flanker distance at a different rate across groups, as shown by the significant interaction between target-flanker distance and group (F_(4,100)_=4.02, p=0.005, ηp^2^=0.13). With respect to the control group, the DD group showed a particularly poor performance in target orientation discrimination (higher IES, reflecting the combination of lower response accuracies and longer response times) when the flankers were shown at the closest distances from the target: post-hoc tests showed that the difference in IES across groups was statistically significant for the target-flanker distances of 0.85° (p=0.005), 1.25° (p=0.03) and 1.45° (p=0.03). With larger target-flanker distances the performance across groups became progressively more comparable, and the IES differences were not significantly different for target-flanker distances of 1.85° (p= 0.13) and of 2.25° (p=0.08). The crowding index (Figure 3D), reflecting the difference in IES between the displays with the smallest and the largest target-flanker distance, was significantly higher in DD compared to the control group (DD group: 384.63 ms ± 213.02 ms, control group: 212 ms ± 95.02 ms, t_(25)_=2.81, p=0.009, Cohen’s *d*=1.04).

In sum, the results of Experiment 3 showed that the enhanced orientation discrimination difficulty in the DD group persisted over several distances between target and flankers compared to controls, as indicated by combined accuracy and reaction time measures.

In order to test for the possibility that the across group differences observed in the orientation discrimination thresholds under crowding (Experiment 2) and in the spatial extent of visual crowding (Experiment 3) were driven by the results of participants having associated reading difficulties or attentional deficits, we repeated the analysis of Experiment 1-3 on a reduced group of eight DD subjects, after excluding those who scored more than 2 standard deviations distance from the control group’s mean in speed or accuracy in the reading or visual search tests.

For the orientation discrimination thresholds measured in Experiment 1 and 2, the interaction between crowding condition and group was significant (F_(1,21)_ = 4.92, p=0.03, ηp^2^=0.19) and the post-hoc tests showed that the increase in orientation discrimination thresholds under the crowded with respect to the uncrowded condition, observed both for the DD (t_(7)_=-4.35, p=0.003, Cohen’s d=1.54) and for the control group (t_(14)_=-9.04, p<10^−5^, Cohen’s d=2.32), was stronger for the DD group with respect to the control group (t_(21)_=2.41, p=0.02, Cohen’s *d*=1.05). Across group differences in orientation discrimination thresholds in the uncrowded condition (Experiment 1), remained not significant (t_(21)_=-1.20, p=0.24, Cohen’s d=0.53).

Reaction times measured in Experiment 3 significantly decreased with target-flanker distance in both groups (significant main effect of target-flanker distance: F_(4,84)_=10.35, p<10^−5^, ηp^2^=0.33), however the DD group was overall slower with respect to the control group (significant main effect of group: F_(1,25)_=9.10, p=0.007, ηp^2^=0.30, significant interaction between group and target–flanker distance: F_(4,84)_=3.08, p=0.007, ηp^2^=0.15 all post-hoc p-values <0.05).

The IES showed a significant interaction between group and target-flanker distance (F_(4,84)_=4.88, p=0.01, ηp^2^=0.18). The IES difference across groups tended to attenuate for larger target-flanker distances, and was significant for most of them - i.e. for target-flankers distances of 0.85° (p=0.001), 1.25° (p=0.01), 1.45° (p=0.005), 1.85° (p= 0.03), but not for 2.25° (p=0.06).

Finally, the crowding index was significantly higher in the DD group with respect to the control group (DD group: 430.58 ms ± 212.30 ms, control group: 212 ms ± 95.02 ms, t_(21)_=3.44 p=0.002, Cohen’s *d*=1.32).

Overall, the analyses performed when excluding from the DD group the four participants with reading and attentional difficulties confirmed the previous results.

### 3.3 Number discrimination

All participants performed two number discrimination tasks in which they were asked to compare the numerosity of two sets of dots.

The DD group was on average less precise with respect to the control group when they had to compare relatively sparse arrays (weber fraction for N24 in the control group=0.17±0.06, in the DD group=0.22±0.06), but not when they had to compare cluttered sets (weber fraction for N64 in the control group=0.17±0.05, in the DD group=0.17±0.06). To statistically test for these differences, we entered the weber fractions into a repeated measures ANOVA with numerosity (2 levels: N24 or N64) and group (2 levels: DD/controls) as within- and between-subject factors, respectively. The ANOVA revealed a significant main effect of numerosity (F_(1,28)_=4.20, p = 0.05, ηp2=0.13), but the main effect of group (F_(1,28)_ = 1.60, p = 0.21 ηp2=0.05), and interaction between group and numerosity (F_(1,28)_= 2.61, p=0.11,ηp2=0.08) were non-significant, suggesting that the two groups were overall less precise in comparing sparser with respect to denser arrays of dots. This latter finding is mainly driven by the DD group that showed a statistically larger weber fraction for comparing sparse (N24) compared to dense (N64) arrays (t_(14)_=2.36 p=0.03), while in the control group this difference was not observed (t_(14)_=0.34 p=0.73). The difference in weber fraction between groups was close to significance for the sparse arrays (t_(28)_=1.90 p=0.06) and not significant for the dense arrays (t(28)=0.17 p=0.86).

### 3.4 Correlation analyses

In order to explore the relation between visual crowding effects, numerosity perception and mathematical abilities, we performed correlation analyses based on Pearson correlation. We correlated measures of crowding that resulted different between groups with numerosity perception and mathematical abilities. Specifically we correlated: 1) the orientation discrimination thresholds under crowding as measured in Experiment 2, 2) the crowding index calculated from Experiment 3, 3) the weber fraction for N24, 4) the weber fraction for N64, 5) the IES score for digit comparison, 6) the IES score for calculation and 7) the IES score for general math ability. Given that some of the DD subjects presented reading and visuo-spatial attention difficulties we performed the correlation analysis with and without regressing out accuracy and speed measured both in the reading and in the visual search tests.

Orientation discrimination thresholds under crowding correlated with IES for digit comparison (r_(27)_=0.56, p=0.002, Fig.4A) but not with IES for calculation (r_(27)_=0.27, p=0.17) nor with IES for general math (r_(27)_=0.28, p=0.14), nor with the weber fraction at N24 (r_(27)_=-0.14, p=0.46) or N64 (r_(27)_= -0.03, p=0.85). The correlation between orientation discrimination thresholds under crowding and IES for digit comparison remained significant even when controlling for reading and visual search abilities (r_(20)_=0.60, p=0.002, Fig. 4B).

**Figure 4.**
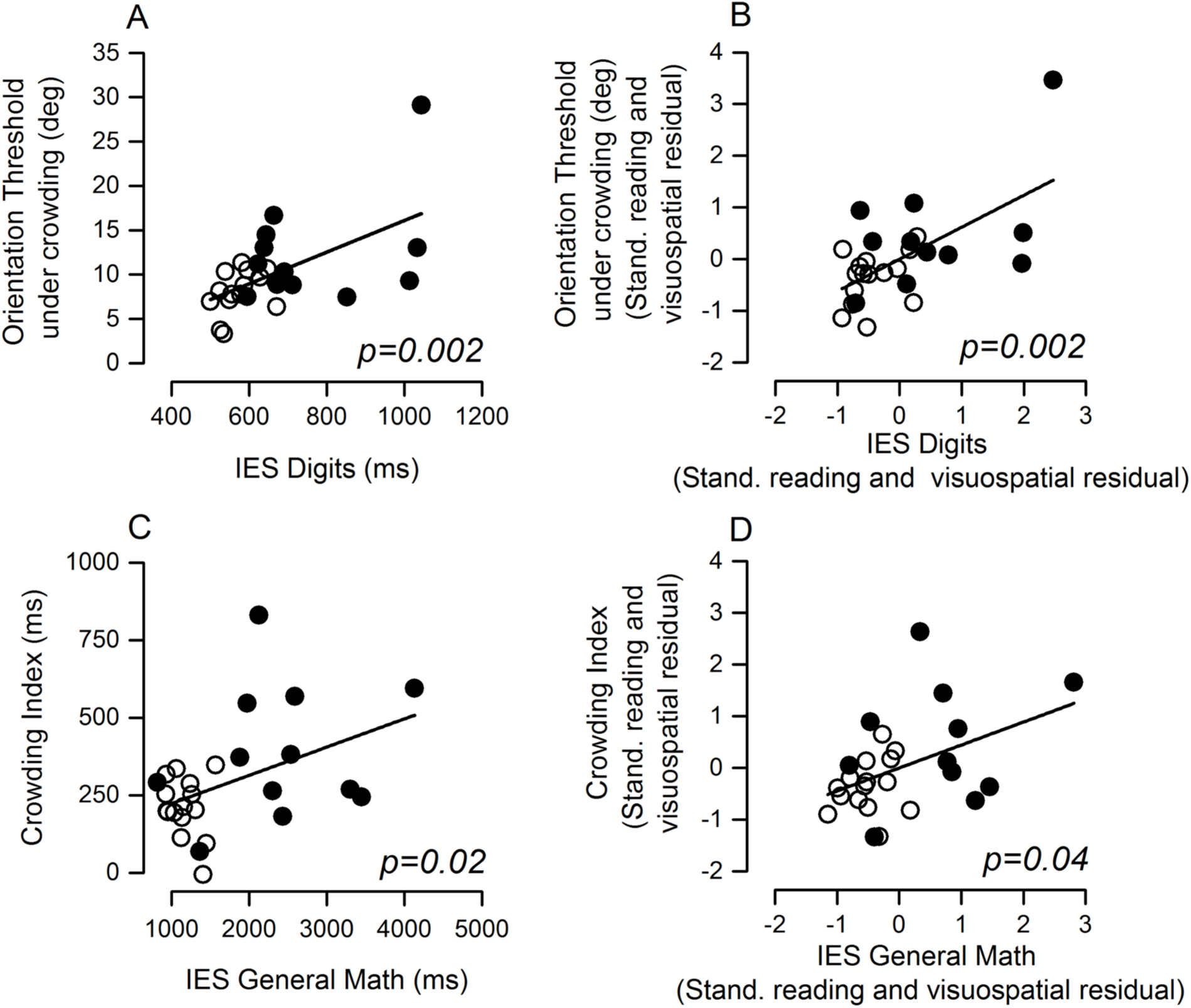
Correlation analysis. Correlation analyses between numerical indices and crowding measures and between their standardized corrected residuals. Orientation discrimination threshold under crowding plotted as a function of IES for digit comparison (A) and plot of the standardized residuals when performance in the reading and visual search tests (B) were regressed out. Same plot but for the crowding index as measured in Experiment 3 (C), and for the standardized residuals (D). White symbols represent control participants, black symbols represent DD participants.

The crowding index correlated with IES for digit comparison (r_(27)_=0.45, p=0.02), with IES for calculation (r_(27)_=0.42, p=0.03) and with IES for general math (r_(27)_=0.45, p=0.02, Fig. 4C), but not with the weber fraction for N24 (r_(27)_=0.09, p=0.64) or N64 (r_(27)_=0.28, p=0.16). The correlations between the crowding index and IES for digit comparison and for general math remained significant after controlling for reading and visuo-spatial attention abilities (r_(20)_=0.54, p=0.009; r_(20)_=0.44, p=0.04, Fig. 4D), while the correlation with IES for calculation was at significance (r_(20)_=0.41, p=0.05).

## 4. Discussion

The aim of the present study was to test whether visual crowding mechanisms are altered in individuals with DD independently of major reading and attentional deficits and whether such an impairment relates to the numerical or arithmetical difficulties. Two groups of participants with and without DD were tested with an orientation discrimination task in different crowding conditions. The orientation discrimination thresholds were comparable across groups when the target grating was presented in isolation. When the target grating was surrounded by flankers, the orientation discrimination thresholds increased in both groups, suggesting that they were both subject to crowding effects. Importantly however, the increase in orientation discrimination threshold under crowding was much higher in the DD group with respect to the control group, pointing at stronger crowding effects.

Increasing the target-flanker distance mitigated the detrimental effect of crowding on the orientation discrimination accuracy in both groups to a comparable extent, yet to perform the task, the DD group needed much longer response time at all the target-flanker distances tested with respect to the control group. This difference could not be accounted for by a general tendency to provide slower responses in the DD group, given that reaction times were comparable across groups when participants were required to perform the orientation discrimination task on the isolated target. Evaluation of participants’ performance in light of the trade-off between accuracy and reaction time, revealed that the excessive crowding effects observed in the DD group compared to control group extended over several larger target-flanker distances. Overall these results point at the presence of enhanced crowding effects in DD that span over a larger than normal spatial extent.

Importantly, the poorer orientation discrimination performance under crowding with respect to the control group was observed even when removing participants with reading or attentional difficulties from the DD group, suggesting that excessive crowding can characterize DD, independently of dyslexia and attentional disorders.

Several models explaining visual crowding phenomena have been proposed (for a recent review see: Manassi and Whitney, 2018). Perceptual failure under crowding has been often attributed to excessive feature integration (Pelli et al., 2004) or information pooling over a large integration area (Parkes et al., 2001). According to this account, crowding occurs when integrating the output of multiple features detectors: target and flankers would fall within the same ‘integration field’ and thus be perceived as jumbled together. In this framework, crowding effects are generally assumed to occur at an early, pre-attentive stage of visual perception. Opposite to this view, the attentional resolution model of crowding proposed that crowding is due to poor resolution of attention (He et al., 1996). According to this account, spatial attention cannot be directed to individual items if they fall within the minimal selection region of attention, in which case items are perceived as a group, preventing their individual identification (Intriligator and Cavanagh, 2001). In line with this theory, some studies found that directing attention to the target location by pre-cueing it improved identification accuracy, reduced critical distance and recognition contrast threshold (Strasburger, 2005; Yeshurun and Rashal, 2010). More recently, the hierarchical sparse selection model proposed that crowding is not due to degraded sensory representations, but to impoverished sampling of such representations by perception (Chaney et al., 2014). In this view, which can successfully explain why crowding occurs at multiple levels of visual analysis, the limiting factor determining crowding would not be the minimal area of the visual field over which attention can operate, but the sparsity of representation sampling within that region.

In light of these models, DD might be associated with either larger integration fields, coarser attentional resolution or reduced ability to sample information. Regardless of the underlying mechanism, what could be the impact of this impairment on the development of numerical and arithmetical cognition?

We hypothesised that due to excessive crowding, during development DD individuals could have more often perceived ensembles as being too cluttered for individual items to be clearly segregated. As a consequence, this could have led to a ‘fuzzy’ representation of numerosity, a less precise association between non-symbolic and symbolic numbers and ultimately less efficient arithmetical abilities.

Independent of the specific origin of visual crowding, this hypothesised chain of events should have resulted in a predictive relationship between the strength of visual crowding effects and both non-symbolic/symbolic number discrimination and calculation abilities. In particular, given that the ability to discriminate the numerosity of sparse, but not of cluttered arrays, was found to be predictive of symbolic mathematical abilities in children, we expected visual crowding to be predictive of numerosity discrimination abilities for sparse arrays. The current results did not support this prediction: we did not find a significant correlation between indices of visual crowding and weber fractions for sparse (nor for dense) dot ensembles. One reason why we might have failed to find such an effect might be related to the fact that we tested adult participants, who might have developed compensatory strategies for dealing with numerosity tasks. For example, participants might have based their decisions on some other visual features, such as the total contrast, density or surface area which in the current experiment increased with numerosity. This might also explain why we only observed a trend for significance when comparing weber fractions for sparse arrays between the DD and the control group. This interpretation fits well with the results reported by previous studies: weber fractions measured with a non-symbolic numerosity discrimination task in DD children differed from controls only when the non-numerical dimensions varied incongruently with the numerical ones (e.g. when the numerically larger set had smaller item surface area compared to the other set) so that it could not be used as a reliable proxy for numerosity (Bugden and Ansari, 2016). Other studies found that numerosity judgments in both DD children (Szűcs et al., 2013) and adults (Castaldi et al., 2018b) were subject to enhanced congruency effects and were strongly biased by non-numerical dimensions (such as, total surface area, edge length, average item size), compared to age-matched control groups. While the current results do not allow us to affirm the link between degree of crowding and non-symbolic numerical acuity, future studies should test for such a link in children with and without DD, as well as under a wider range of controls for non-numerical quantitative properties.

It is also important to note that the role of non-symbolic numerical abilities in the development of integer concepts has been challenged by some authors (Carey and Barner, 2019) and that the predictive relationship between non-symbolic number discrimination and formal arithmetic has sometimes not been replicated. Rather, symbolic number abilities resulted to be a more robust predictor of arithmetical skills across studies (De Smedt et al., 2013; Schneider et al., 2017). In the current study we observed that the orientation discrimination thresholds under crowding and the crowding index were predictive of the performance in both a digit comparison task and general math skills: participants with stronger visual crowding effects performed more poorly in digit comparison and calculation tasks. Given that in the digit comparison task, two numbers were presented far apart from each other, on the two sides of the screen, it is very unlikely that they would have been displayed within the same ‘integration field’.

In the current study we ruled out the possibility that the observed across group differences in crowding effects were exclusively driven by the participants with major visuo-spatial attentional deficits, potentially identifying participants with a history of ADHD. Yet, it remains possible that some attentional weaknesses, known to be present in pure DD individuals (Askenazi and Henik, 2010), although potentially not strong enough to result significantly different with respect to the control group when measured with the visual search test used here, might have contributed to the excessive crowding effects observed. Related to this interpretation, another hypothesis that can be advanced to explain the observed relation between crowding and numerical abilities is that attentional weaknesses might have prevented the development of a sufficiently clear spatial representation of numbers. An influential hypothesis proposed that numbers are internally represented along a spatially oriented number line (Dehaene et al., 1993; Dehaene, 2003). According to this view, spatial associations are relevant both for understanding the meaning of numerical values, which would be conveyed by their position on the number line, as well as during calculation, which is thought to involve shifts of spatial attention along the number line (Hubbard et al., 2005). There is evidence that impairments of visuo-spatial attention affect this numerical space: neuropsychological studies on patients with hemi-spatial representational neglect found that they perform extremely poorly in number bisection tasks, most likely because the deficit in orienting visuo-spatial attention to the contralesional hemispace also extended to the internal representation of the number line (Zorzi et al., 2002, 2006). If the excessive crowding effects observed in the current study in the DD group reflect a limited spatial resolution of attention, then they might have hampered the development of a clear internal representation of numbers in space. Numbers might be ‘too crowded’ along the internal mental number line to be sharply sampled and manipulated during number comparison or calculation tasks. The possibility that crowding may degrade internal representations, and not only visual percepts, is also supported by a recent study showing that crowding also affects visual working memory contents (Tamber-Rosenau et al., 2015). Such ‘representational crowding’ might set limits to the ability of the DD individuals to precisely select the two numerical values to be compared on the number line and/or to keep them in memory during the comparison or calculation process.

It has been previously suggested by some authors that the difficulties in calculation observed in DD might be more related to impaired general executive functions such as attention, working/long-term memory and inhibition rather than to conceptual knowledge of number and arithmetic (Bull and Scerif, 2001; Geary, 2004). The relation between excessive crowding effects and numerical and calculation abilities observed in the current study could be interpreted as an evidence in support of that model of DD, with no need to evoke any domain specificity. However, arguing against such a pure domain-general hypothesis, it has been shown that training general attention in DD adults (using video-games) improved the orienting system, but did not improve the difficulties in the DD group in arithmetic or in numerical processing (Ashkenazi and Henik, 2012). Neither did specifically boosting the alerting system with brief auditory cues prior to an estimation task increase the smaller than normal subitizing range or accuracy in DD individuals (Gliksman and Henik, 2019). Moreover, domain-general deficits in DD are sometimes reported to be specific to the numerical stimuli. For example, one study found that verbal working memory deficits in DD were stronger when digits were tested with respect to letters or words (Peng and Fuchs, 2016). Others found that inhibition deficits in DD appeared only with Stroop paradigms involving digits, but not letters or geometric features (Wang et al., 2012). In ensemble discrimination tasks, enhanced interference from the task-irrelevant dimensions manifested only when DD subjects were required to judge the numerosities of the ensemble, and not when judging other quantitative features, such as its average item size (Castaldi et al., 2018b). Overall, this evidence suggests that, although DD may present some weakness in domain-general functions, these may not uniquely explain the specific numerical difficulties in DD. Rather it is more likely that weak domain-general functions might interact with an impaired number specific magnitude system. Using digit stimuli, some studies found that flankers might influence target detection not only because of their perceptual similarity, but also because of their semantic closeness: responses are facilitated when a target number is surrounded by numerically congruent flankers and asymmetries in flanker-target interference occur when magnitude or parity of the target (Huckauf et al., 2008), but not its physical characteristics, need to be extracted (Patro and Huckauf, 2019). To compare domain-general vs domain-specific attentional deficits in DD, future studies should test whether DD individuals present even stronger crowding effects when using numerical with respect to non-numerical stimuli (such as the Gabors used here). Moreover, in order to test our hypothesis that crowding impacts numerical cognition by preventing the development of a clear internal number line, it would be interesting to test whether pre-cueing attention to the operands’ locations along the number line would improve performance in calculation tasks by increasing the ‘mental critical spacing’ between numbers.

In conclusion, this study provides a first report of abnormal crowding in DD individuals, which can be found independently of pronounced associated reading and visual-attention deficits. Which are the exact mechanisms underlying the excessive crowding effects and how they contribute to the development of numerical and arithmetical difficulties in DD needs to be explored further in future studies. Perhaps, similarly to what was previously observed in dyslexic individuals (Martelli et al., 2009), excessive crowding may contribute to create numerical difficulties in DD, although not fully accounting for it.

## Acknowledgements

This research has received funding from the French National Research Agency (grant No ANR-14-CE13-0020-01 to E. Eger), from the European Research Council (ERC) under the European Union’s Horizon 2020 research and innovation programmes (grant agreement No 801715 – PUPILTRAITS) and from the Accademia dei Lincei (fellowship “G. Guelfi per le ricerche nel campo della biomedicina o della biologia 2019”).

